# Freddie: Annotation-independent Detection and Discovery of Transcriptomic Alternative Splicing Isoforms

**DOI:** 10.1101/2021.01.20.427493

**Authors:** Baraa Orabi, Brian McConeghy, Cedric Chauve, Faraz Hach

## Abstract

Alternative splicing (AS) is an important mechanism in the development of many cancers, as novel or aberrant AS patterns play an important role as an independent onco-driver. In addition, cancer-specific AS is potentially an effective target of personalized cancer therapeutics. However, detecting AS events remains a challenging task, especially if these AS events are not pre-annotated. This is exacerbated by the fact that existing transcriptome annotation databases are far from being comprehensive, especially with regard to cancer-specific AS. Additionally, traditional sequencing technologies are severely limited by the short length of the generated reads, that rarely spans more than a single splice junction site. Given these challenges, transcriptomic long-read (LR) sequencing presents a promising potential for the detection and discovery of AS.

We present Freddie, a computational annotation-independent isoform discovery and detection tool. Freddie takes as input transcriptomic LR sequencing of a sample and computes a set of isoforms for the given sample. Freddie takes as input the genomic alignment of the transcriptomic LRs generated by a splice aligner. It then partitions the reads to sets that can be processed independently and in parallel. For each partition, Freddie segments the genomic alignment of the reads into canonical exon segments. The goal of this segmentation is to be able to represent any potential isoform as a subset of these canonical exons. This segmentation is formulated as an optimization problem and is solved with a Dynamic Programming algorithm. Then, Freddie reconstructs the isoforms by jointly clustering and error-correcting the reads using the canonical segmentation as a succinct representation. The clustering and error-correcting step is formulated as an optimization problem – the Minimum Error Clustering into Isoforms (MErCi) problem – and is solved using Integer Linear Programming (ILP).

We compare the performance of Freddie on simulated datasets with other isoform detection tools with varying dependence on annotation databases. We show that Freddie outperforms the other tools in its recall, including those given the complete ground truth annotation. In terms of false positive rate, Freddie performs comparably to the other tools. We also run Freddie on a transcriptomic LR dataset generated in-house from a prostate cancer cell line. Freddie detects a potentially novel Androgen Receptor isoform that includes novel intron retention. We cross-validate this novel intron retention using orthogonal publicly available short-read RNA-seq datasets.

**Availability:** Freddie is open source and available at https://bitbucket.org/baraaorabi/freddie

## 1 Introduction

Alternative splicing (AS) is a cellular process that enables a single gene to encode for multiple different proteins [4], contributing to an increased protein diversity [2, 22]. Recent findings show that AS plays a critical role in regulating gene expression [9] and tissue specialization [32]. Abnormalities in the AS of genes are also linked to the pathogenesis of many diseases, including cancer. For example, AS aberrations contribute to the tumour’s ability to proliferate and to evade programmed cell death [24]. Additionally, cancer-specific AS aberrations present potential targets for cancer therapeutics [13, 6]. The efficacy of these AS-centered treatments relies on the personalized, accurate, and rapid detection of AS isoforms at the level of individual patients.

From the mid-2000s and until recently, short-read (SR) sequencing has been the dominant high-throughput sequencing technology for genomics and transcriptomics analysis. While SR sequencing has a very low sequencing error rate (of the order of 1 error per 1000 sequenced nucleotides), it generates reads that are too short to fully sequence an isoform molecule in a single read: 75-250 nucleotides per read vs a median of 909 nucleotides per isoform [7, 35]. SR AS detection methods use SRs to identify splicing junctions between pairs of exons [26, 31, 17, 16]. However, due to the limited span of SRs, such methods face major challenges in “chaining” these splicing junctions to reconstruct complete isoforms.

More recently, long-read (LR) sequencing became a commercially viable option to study the transcriptome and genome [1]. In theory, LR sequencing machines can sequence the full length of RNA molecules generating reads that range from thousands to tens of thousands of nucleotides [5]. Thus, ideally, aligning LRs to the reference genome should be enough to perfectly define the exons of the underlying isoforms. However, in practice, LRs suffer from a high sequencing error rate (10-20 errors per 100 sequenced nucleotides) that is dominated by indels (erroneous insertions and deletions) [11]. When comparing aligned LRs to a reference genome, indels cause erroneous shifts in the observed boundaries of the exons and occasionally result in missing smaller exons altogether. Additionally, LR sequencing of an RNA molecule, more often than not, terminates before reaching the full length of the molecule, resulting in missing exons at the tail of the isoform [28]. These main types of LR sequencing errors are summarized in Figure S1. Current isoforms detection methods based on LRs overcome these challenges by relying on existing isoform annotation databases. The current methods can be classified into three categories depending on how they use available annotation data: (i) methods that detect only isoforms that are described in the annotation which is typically done by aligning the LRs to the sequences of annotated isoforms; (ii) methods that detect isoforms with potentially novel exon chains if those exons boundaries are present in some annotated isoforms (e.g FLAIR [30]); (iii) de-novo methods that do not rely on any annotation but can potentially benefit from using existing annotation data (e.g. StringTie2 [12]). Approaches that rely on annotation data are limited by the fact that annotation databases are incomplete: millions of splicing genomic locations and thousands of isoforms are present in different individuals but not in major annotation databases [21, 34]. Figure 1 provides further detail into the hierarchy of dependence on annotations for isoform detection.

**Fig. 1.**
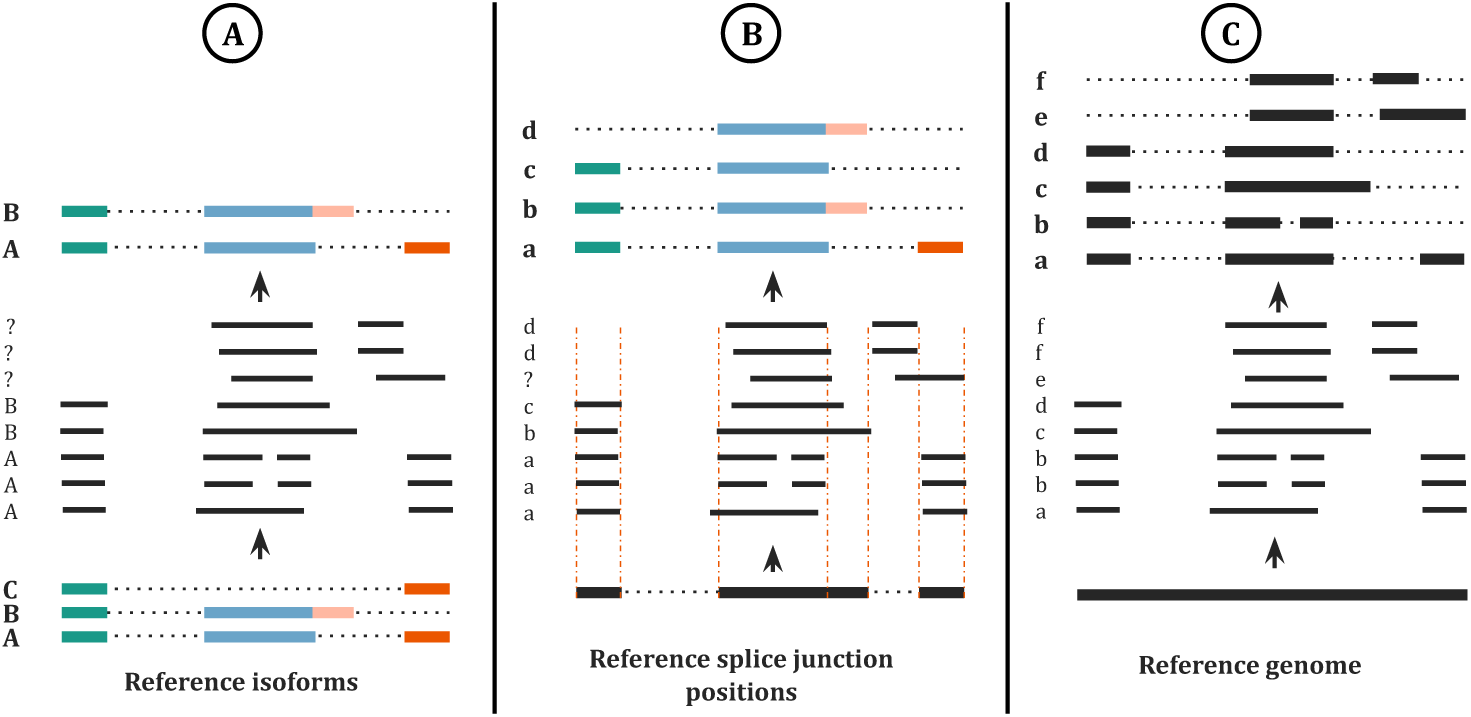
AS isoform detection tools can be put on a spectrum in terms of their reliance on reference annotations. A) On the left of the spectrum, the tool is fully dependant on the known isoform annotations and is thus unable to discover any novel isoforms. Each read is annotated by the isoform that it best matches or is discarded if it does not match any isoform. B) In the middle, the tool is partially reliant on the isoform annotation; novel isoforms can be detected as long as they are composed of known splice junctions (i.e. boundaries of known isoforms). The split-alignment boundary of each read is corrected to best matching splice junction position. If a read has a novel splice junction position, the tool will have difficulty identifying its isoform structure. C) On the right, the tool does not rely on the reference annotation. Instead, it relies solely on the split-alignment of the reads to the reference genome.

### Contributions

In this paper, we introduce Freddie, a multi-stage novel computational method aimed at detecting isoforms using LR sequencing without relying on isoform annotation data. The design of each stage in Freddie is motivated by the specific challenges of annotation-free isoform detection from noisy LRs. We compare the accuracy of Freddie against two alternative state-of-the-art isoform detection tools, FLAIR [30] and StringTie2 [12]. We show using simulated data that Freddie achieves accuracy on par with FLAIR despite not using any annotations and outperforms StringTie2 in accuracy. Furthermore, Freddie’s accuracy outpaces FLAIR’s when FLAIR is provided with partial annotations. Finally, we demonstrate Freddie’s ability to detect novel isoforms on a real cancer cell line dataset.

## 2 Methods

Freddie takes as input the mapping of transcriptomic long reads to a reference genome and outputs, in GTF format, a list of detected isoforms each described as a set of genomic intervals. Freddie assumes that the mapping is performed by a splice-aware mapper which attempts to solve the problem of finding the best read-to-genome alignment that accounts for introns by allowing for large deletions to not be penalized. By default, we use deSALT [19] mapper since it is designed for split-aligning LRs and does not require any annotation about the expected splice-junction sites. Freddie is composed of three stages: partitioning, segmentation, and clustering/error-correction as depicted in Figure 2. The design of each of these stages is motivated by the challenges of isoform detection using noisy long-read sequencing that does not rely on annotation data.

**Fig. 2.**
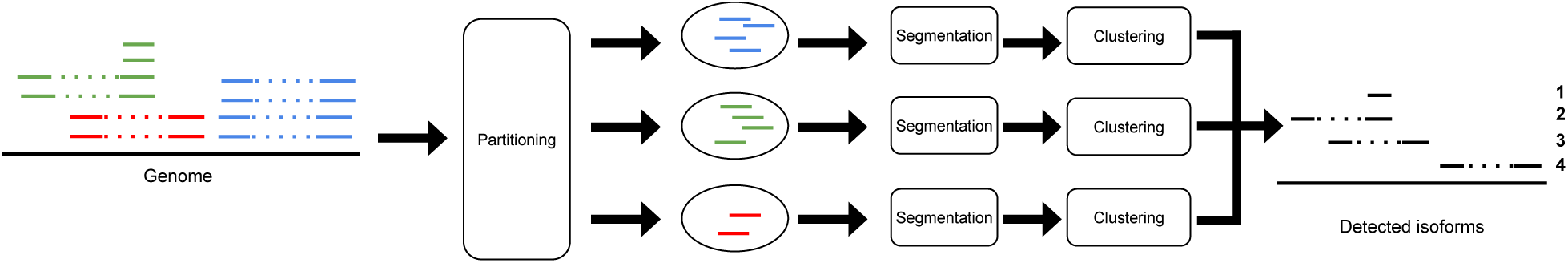
Overview of the Freddie stages, from genomic split-alignments of LRs to detected isoforms.

### 2.1 Read partitioning

In this stage, we partition the aligned reads into sets with the aim that no isoform from the sequenced transcriptome has reads present in different sets. As a result, each set of the partition can be assumed to contain all the reads from a set of isoforms and can be processed independently from the other sets, which allows for highly parallel processing. We define the partition sets as follows: 1) if two reads have split-alignment intervals that overlap, then they must be in the same set; and 2) if two reads are in different sets, then their split-alignment intervals cannot overlap.

Given a sorted list of the split-alignment intervals of the reads, we can compute the partition sets in linear time. This is achieved by first identifying genomic intervals with contiguous coverage, and then running a classic breadth-first search using the read membership in these contiguous genomic intervals. At the end of this stage, we have independent sets of reads that we can process independently and in parallel in the next stages. Therefore, when we refer to the reads in the next stages, we mean only the reads in a given set. Figure 2 illustrates the parallel processing of multiple partition read sets in the next stages.

### 2.2 Canonical segmentation of the genome

The segmentation stage addresses the challenge of detecting exon (and intron) boundaries from the alignments of LRs to a reference genome. Annotation-dependent isoform detection tools bypass this challenge by matching the LR alignments to the closest exon boundary in the annotation, assuming that any deviation from the annotation is a result of sequencing noise and not of a potentially novel AS event. To overcome this limitation, we propose a data-driven segmentation approach aiming at identifying exon boundaries by finding a set of segmentation breakpoints that are best supported by the input LR split-alignments. To find this segmentation, we devise a two-step process (illustrated in Figure 3).

**Fig. 3.**
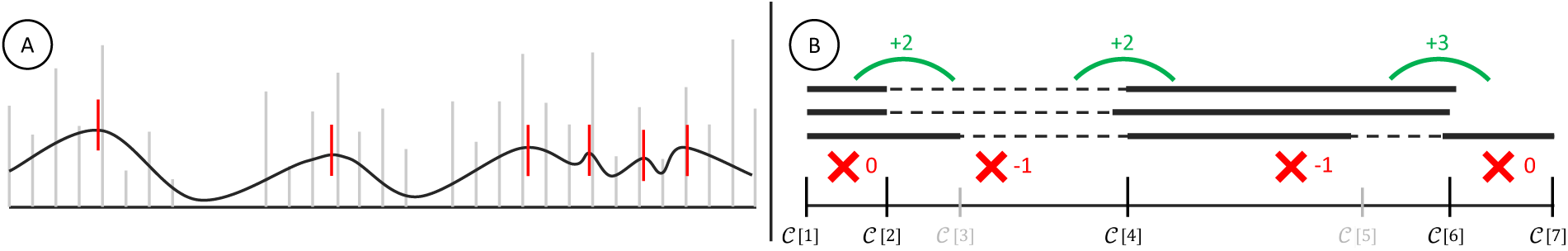
Illustrating the two main steps of Freddie’s segmentation stage. A) The interval boundaries of the genomic split-alignments of the reads are represented as a discrete signal (gray). This signal is smoothed using a Gaussian filter. The peaks (red) of this filtered signal are used as candidate breakpoints for the segmentation. B) A subset of the elements in the list of candidate breakpoints, *𝒞*, is selected and scored. Score increases due to high coverage contrast are shown in green and score decreases due to partial coverage are shown in red. This scoring scheme is used to select the optimal subset of *𝒞* to be used as canonical segmentation in this stage.

#### Identifying candidate breakpoints

To generate the set of candidate breakpoints, *C*, we treat the LR split-alignments as a discrete signal: for a genomic position *i*, we define *M* [*i*] as the number of reads that have a split-alignment interval starting or ending on *i*. We then apply a Gaussian filter on the signal encoded by the array *M* to smooth over the signal’s noise. The Gaussian filter smooths out the raw signal and makes it more robust to the noise due to indel sequencing errors on the potential splicing positions. We denote the smoothed signal by ℳ. Finally, we denote by *C* the set of genomic positions of the local peaks (i.e. local maxima) of ℳ (see Figure 3A). Our implementation uses scikit-learn [25] Gaussian filter and peak finding functions.

#### Pruning the set of the candidate breakpoints

Freddie selects a subset of breakpoints, *S* ⊆*C*, to represent the finalized set of canonical exon boundaries inferred from the input long reads alignments. Note that the breakpoint sets, *C* and *S*, divide the genome, respectively, into |*C* |−1 and |*S* |−1 non-overlapping genomic segments. We define a scoring function, *f*, which given a set of breakpoints simultaneously rewards individual reads for sharp changes in coverage between consecutive segments and penalizes them for having partial alignment on individual segments (see Figure 3B). Freddie selects the subset *S* which maximizes *f* (*S*) over all the subsets of *C*, using a Dynamic Programming (DP) algorithm.

To formally define *f* (*S*), we introduce the following notations. Let *cov*(*r, i, j*) be the percentage of positions in genomic segment [*i, j*] that are covered by the split-alignment intervals of read *r*. We also define three binary indicators, *FC_r,i,j_, NC_r,i,j_* and *PC_r,i,j_*, to respectively indicate if read *r* is covering, not covering, or partially covering the genomic segment [*i, j*]:

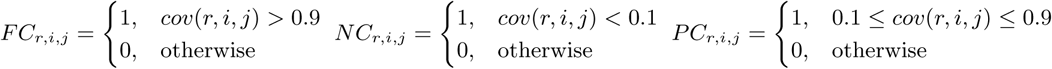

Next, we define the number of reads with partial split-alignment coverage over a genomic segment [*i, j*]:

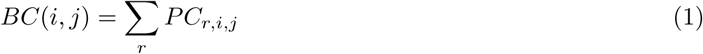

Finally, the number of reads with sharply contrasting coverage between two neighboring genomic segments, [*i, j*] and [*j, k*], *GC*(*i, j, k*), is defined as follows:

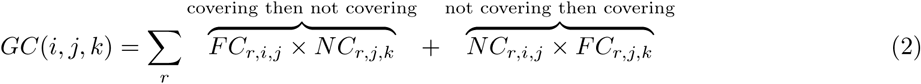

Using these functions, we define the scoring function *f* :

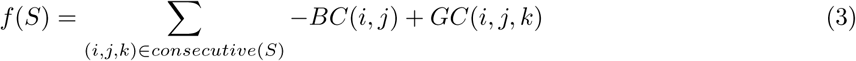

Where *consecutive*(*S*) is the set of all triplets of consecutive breakpoints in *S*.

We want to find *S* which maximizes *f* (*S*) from all possible subsets of *C*. Although there are exponentially many subsets of *C*, we can efficiently find the optimal subset *S* because the structure of the scoring function lends itself readily to cubic time DP. To describe the DP equations, let C be the list of the elements in *C* sorted by increasing genomic coordinate and let *𝓁* = |𝒞|. We denote by 𝒞 [*i*] the breakpoint at index *i* in 𝒞 and by 𝒞 [*i..j*] the sub-list of C starting at index *i* and ending at index *j* (inclusively on both ends). We fill in a DP table *D* defined as follows: for 1 ≤ *h < m < n* ≤ *𝓁, D*[*h, m, n*] is the optimal score (as defined in Equation 3) among all subsets of *C* composed of the breakpoints in 𝒞 [*h..𝓁*] with the additional constraint that the first three breakpoints in the subset, according to the order defined by 𝒞, occur at the respective indices *h, m* and *n* in 𝒞. Then:

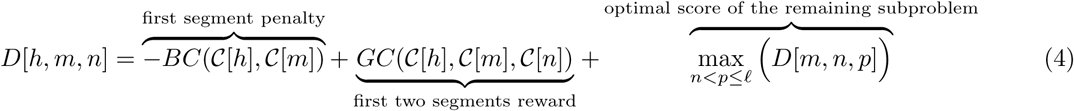

The score of the optimal segmentation *S* is then:

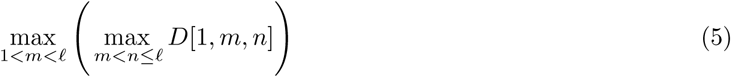

Finally, to obtain *S*, we simply backtrack through the DP table *D*.

#### Speed-up heuristics

Since the cubic time complexity can be prohibitive if there is a large number of candidate breakpoints, we reduce the complexity by fixing breakpoints using some heuristics before running the DP formulation. Fixing breakpoints effectively breaks down the problem into independent subproblems, reducing the base of the cubic time complexity and thus dramatically speeding up the effective runtime. The first heuristics we use is to select any breakpoint that has extremely high smoothed signal value since it is highly likely to be in the optimal set of breakpoints. Specifically, we fix breakpoints with values higher than the mean plus three times the standard deviation of the ℳ[*i*] for all *i* in *C*. The second heuristics we use is to introduce a limit on the maximum allowed problem size. If the size of *C* after the first heuristic exceeds this limit, then we split it uniformly into subproblems, each of size less than or equal to the limit. The limit is a user-set parameter with a default value of 50.

#### Vectorization of LRs

For the next stage, we use the selected breakpoints, *S*, of this stage to succinctly represent the reads: each LR is encoded as a binary vector of length *V* = | *S* | − 1 in which each bit indicates the presence (1) or absence (0) of a segment on the LR using the 90% coverage threshold used in above, as illustrated in Figure 4.

**Fig. 4.**
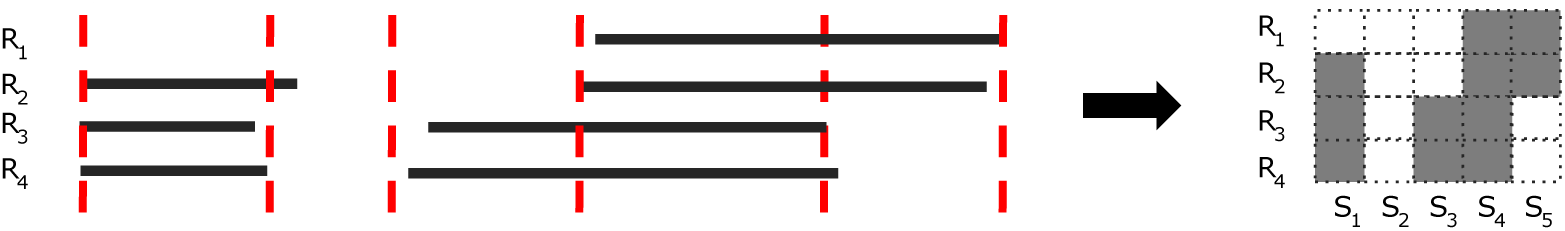
The segmentation is projected on the reads. Each read is represented as a binary vector of size equal to the number of canonical segments. A segment has a value of 1 (filled gray) if a read has at least 90% coverage over the segment, and 0 otherwise.

### 2.3 Clustering and error correction

The goal of this stage is to compute a set of potential isoforms using the succinct binary vector representation of the LRs generated by the segmentation stage. Our approach is to cluster the LRs such that each cluster represents a potential isoform. A crucial point is to consider the possibility for some reads to have erroneously missed segments (1 to 0 errors) due to sequencing errors and to correct such errors by using evidence from other LRs from the same cluster. Thus, each cluster is composed of similar reads that potentially originate from the same isoform. We can then reconstruct each isoform by generating the consensus structure from the vector representation of the LRs in the cluster. We recognize two main challenges for this task that are not faced by annotation-dependent tools: we do not know a priori (i) the number of clusters (i.e. isoforms) and (ii) the structure of the isoforms in terms of segments. To overcome these challenges, we devise an iterative process inspired by the Minimum Error Correction (MEC) problem [18] that is commonly used in haplotype assembly.

In each round of this iterative process, we assign the input reads into one of two bins: an isoform bin and a recycling bin. The isoform bin should contain similar reads that are assumed to originate from the same isoform, while the recycling bin should contain everything else. At the end of each round, the isoform bin reads are set aside and their consensus is used as the structure of a detected isoform, while the recycling bin reads are used as input to the next round (see Figure 5). The assignment of the reads in each round to one of the two bins is done according to a scoring function: each read in the isoform bin incurs a penalty proportional to the number error corrections it requires in order to match the consensus structure of the isoform bin reads (similar to the MEC problem) while each read in the recycling bin incurs a constant penalty. The interesting feature of this approach lies in its ability to cluster the reads without specifying the number of expected clusters or their structure. We call this clustering problem the Minimum Error Clustering into Isoforms (MErCI) problem.

**Fig. 5.**
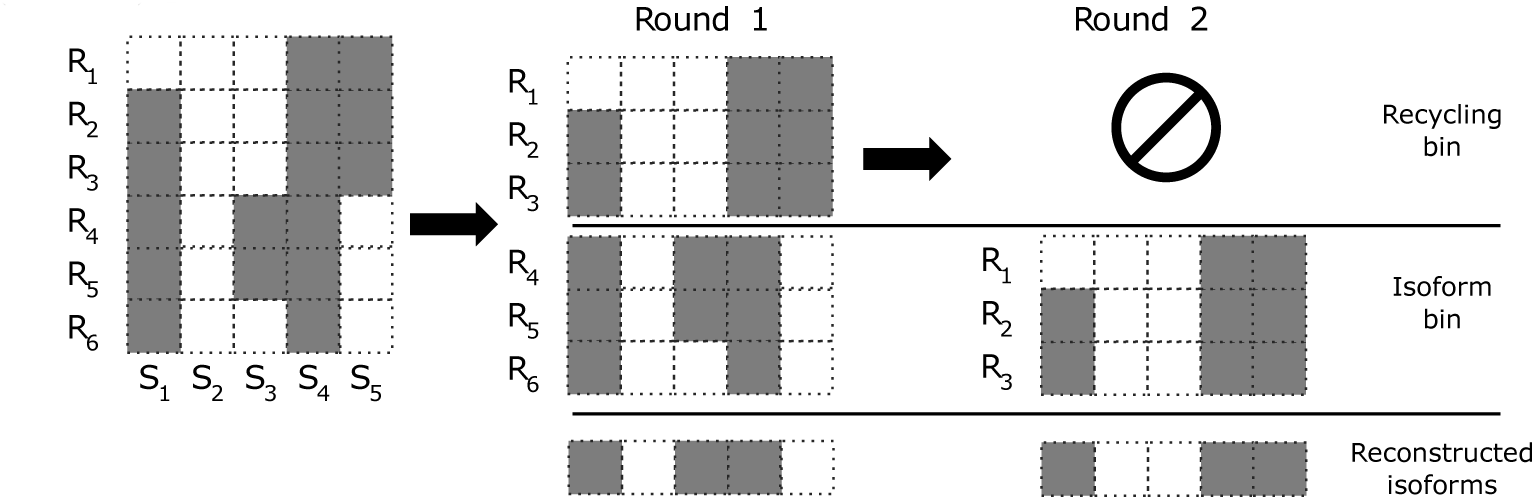
The input to the clustering stage is the vectorized reads. In each round of MErCI, the reads are assigned to either the isoform bin or the recycling bin. The isoform bin reads are set aside and the recycling bin reads are used as input to the next stage. The read vectors of each isoform cluster are used to reconstruct the isoforms using a simple column-wise consensus strategy.

#### The Minimum Error Clustering into Isoforms (MErCI) problem

To formulate the MErCI problem, we start by introducing some preliminary definitions. Let *A* be an *N*×*V* binary matrix representing the input reads, i.e., *N* is the number of reads and *V* = |*S*| − 1 is the number of canonical genomic segments: *A*[*i*][*j*] = 1 if the *i*-th read covers *>* 90% of the *j*-th segment and *A*[*i*][*j*] = 0 otherwise. For a given subset *I* of the reads, we define the *induced isoform* of *I*, ℐ*_I_*, to be a binary vector of size *V* consisting of column-wise disjunction of the reads in *I*:

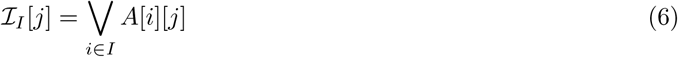

Given an induced isoform, ℐ*_I_*, we define the correction cost relative to ℐ*_I_* of read *i* ∈ *I<, CC*(*i,* ℐ*_I_*), to be the number segments present in ℐ*_I_* but not in *A*[*i*]:

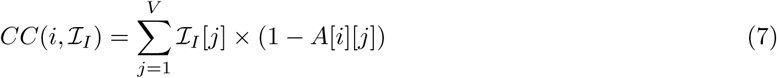

Finally, we define the optimization goal of MErCI to be finding a subset of reads *I* that minimizes:

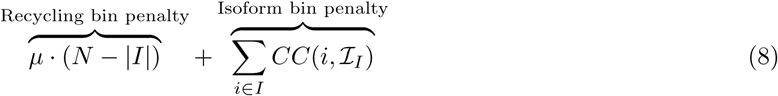

where *µ* is a user-defined threshold forbidding the inclusion of a read into the isoform bin if at least *µ*corrections are required. By default, we set *µ* = 3, allowing for up to two error corrections per read.

#### Additional considerations for transcriptomic LRs

In Freddie, we extend and modify the MErCI problem as described in the previous section in order to include considerations that are specific to transcriptomic LR sequencing data.

#### Poly-A tail

During the biological process of transcription, the spliced RNA molecule is extended with a small sequence of A nucleotides, known as the poly-A tail. The presence of the poly-A sequence is used as a target in the preparation step of sequencing to extract and isolate the mRNA molecules from the sample RNA material. Note that the poly-A tail is not part of the genomic sequence of the gene. Therefore, if a LR has successfully sequenced the poly-A tail of its isoform, we expect to observe an A-enriched sequence (or a T-enriched sequence for cDNA sequencing) in the LR in the parts of its sequence that did not align to the genome. If we observe the A-enriched sequence after the last covered canonical segment of the LR, then we can infer that the gene of the LR isoform is on the forward strand of the genome and we describe this LR as a *forward* read. Similarly, if we observe the A-enriched sequence before the first covered canonical segment of the LR, we describe the LR as a *reverse* read. Note that the order of the segments is defined by their increasing genomic coordinates. In Freddie, we constrain the MErCI assignment to the isoform cluster to forbid assigning a forward read and a reverse read to the isoform cluster in a given round.

#### Truncated reads

As mentioned earlier, LRs do not always cover the full length of an RNA/cDNA molecule. Therefore, for a given LR, it makes sense not to penalize the corrected segments at the start or end of the isoform if the sequencing has terminated before they were sequenced. We, therefore, modify the correction cost function in Freddie to penalize only the internal segments of a given LR. We define the internal segments of a LR to be the segments between the first and last covered segments of the LR (i.e. segments between the first and last 1 entries in the binary representation of the read). For forward reads, we extend the definition of internal segments to include segments after its last covered segment. And similarly, for reverse reads we include segments before its first covered segment. Let *Internal* (*i, j*) = 1 if segment *j* is internal for read *i*, and 0 otherwise. We then redefine the correction cost function to be:

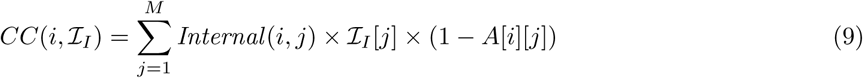

#### Length of corrected segments

When correcting internal segments in a LR, we also take into account the lengths of the corrected segments. This is because missing shorter segments is more likely to be the result of sequencing and mapping errors. More specifically, we want to account for the possibility that the missing segments were sequenced by the LR, but due to substitution and indel errors, the LR alignment to the reference genome missed some segments. Therefore, for each read we extract the set of its internal contiguous missing segments and their flanking covered segments (i.e. maximal stretches of 0’s surrounded by 1 on both side) which we call “gaps”. Formally, we denote a gap with a triplet (*s, e, l*) where *s* and *e* are the first and last covered (i.e. 1) segments of the gap, and *l* is the number of unaligned bases on the read between the *s* and *e*. Then, for each gap (*s, e, l*) of read *i* that is assigned to the isoform bin *I*, we add a restriction that the genomic length of the corrected segments between *s* and *e* must be close to *l*. More precisely, let *CGL*(*s, e, i*) be the genomic length of the corrected segments within the gap (*s, e, l*) of read *i*:

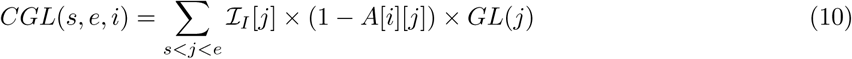

where *GL*(*j*) denotes the genomic length of the *j*-th segment. Then the following length condition must hold:

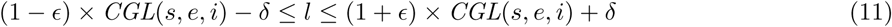

By default, we set *1* to 0.1 and *δ* to 20 nucleotides.

#### Solving MErCI using Integer Linear Programming

The search space for assigning reads to the isoform bin or the recycling bin is exponentially large in the size of the input. To overcome this challenge, we formulate the MErCI problem with the extra constraints described in Section 2.3 as an Integer Linear Program (ILP). We use the ILP solver Gurobi [8] to obtain an optimal assignment of the reads according to the MErCI formulation.

#### Isoform reconstruction

Finally, to reconstruct the isoforms, we build a consensus for each cluster of reads. Here, we apply a simple rule of column-wise consensus of the reads with a plurality threshold of 30%. We chose this low threshold to take into account that fewer reads might successfully sequence the full length of the isoform.

## 3 Results

To evaluate the accuracy and predictive power of Freddie, we assessed Freddie against two popular methods, FLAIR [30] and StringTie2 [12], on a simulated dataset. As the ground truth about the real datasets is not known a priori, it is essential to use controlled simulated data to perform accuracy measurements. However, the simulation is able to only reflect the aspects of LR sequencing that we are aware of and that we introduce in the simulation’s design. Therefore, it is also equally important to test our method on real data set and attempt to validate it using orthogonal technologies to provide some assurance of the usability of our method. Thus, we also tested Freddie on a LR dataset of 22Rv1, a prostate cancer cell line, sample.

### 3.1 Simulated data experiment

#### Data generation

We generated a LR transcriptomic sequencing dataset for chromosome 21 of the human genome. For this dataset, we wanted it to reflect the isoform expression distribution of a typical real dataset. Therefore, we downloaded a real transcriptomic LR sequencing dataset (sequenced on MinION) from the Oxford Nanopore Whole Genome Sequencing consortium [34]. We then mapped the LRs of this real dataset using Minimap2 [15] to the set of annotated human isoform sequences from ENSEMBL 97 database [3] and used the number of primary alignments of the reads to each isoform as an expression profile of this real dataset. We used this expression profile to simulate LRs from the isoforms sequences of chromosome 21 using Badread [33], a simulator specifically designed for Oxford Nanopore LRs. Finally, we discarded any isoform (and its simulated reads) with less than three simulated reads. In total, the simulated dataset includes 26,731 reads from 702 isoforms with a median length of 1,905 nucleotides. The median number of reads simulated per isoform is nine.

#### Tool configuration

We used Minimap2 [15] and deSALT [19] to generate input genomic split-alignments and ran them without using any annotation files. The accuracy performance was very stable for Freddie and FLAIR regardless of the aligner while StringTie2 had a much worse accuracy with using deSALT compared to using Minimap2. To avoid any disadvantage caused by the choice of the aligner, we present Minimap2 results for FLAIR and StringTie2 and deSALT results for Freddie, which gives the best accuracy result for each tool. The complete set of results are available in Figure S2 and S3 of the Supplementary Material. We tested all tools using their default settings. We ran StringTie2 (with -L flag) and Freddie without any annotation files. FLAIR requires the use of an annotation file so we supplied it with the complete GTF file for chromosome 21, including the annotations of the 702 considered isoforms. Note that this provides the best-case scenario for FLAIR in terms of the comprehensiveness of the annotation. To further investigate FLAIR’s reliance on the annotation, we tested it with subsets of the annotation with varying sampling rates of 75%, 50%, 25%, and 1% of the annotation isoforms. We present here the accuracy results of FLAIR using 100% and 50% annotation sampling rates. The complete results for FLAIR are available in Figure S4 of the Supplementary Material.

#### Accuracy measurement

In assessing the accuracy of AS detection, the simulated isoforms should not be treated as independent data points; the structures of isoforms from the same gene can greatly overlap which makes it harder to interpret the classical notions of precision and recall, and similarity between isoforms, either ground truth isoforms or predicted isoforms, should be accounted for. To achieve that, we construct a graph *G* whose vertices are *V* = *P_T_* ∪ *S* where *P_T_* and *S* are, respectively, the sets of isoforms predicted by tool *T* and the set of true isoforms. We add an edge between any pair of isoforms, *i*_1_ and *i*_2_, if their pairwise sequence split-alignment score is higher than a given threshold *t*∈[0, 1]. We define the pairwise sequence split-alignment score to be:where *aln*(*i*_1_*, i*_2_) is the total lengths of the split-alignment intervals of aligning the sequences of *i*_1_ and *i*_2_ isoforms

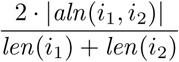

where |*aln*(*i*1,*i*2) | is the length of the sequence of isoform *i*. We then extract all the (connected) components of the graph *G* and classify them into three categories: (i) a mixed clique contains at least one predicted isoform (PI) and at least one ground truth isoform (GTI) and an edge between all pairs of its vertices; (ii) a non-mixed component contains only PIs or only GTIs and may or may not be a clique; and (iii) a mixed ambiguous component contains at least one PI and at least one GTI, with some pair of vertices having no edge connecting them. For a given alignment score threshold, mixed cliques (category i) represent sets of pairwise similar PIs and GTIs that we interpret as true positives, while PIs and GTIs in non-mixed components (category ii) can be interpreted as false positive and false negative, respectively. The isoforms in mixed ambiguous components (category iii) cannot be unambiguously labeled as true positive, false positive or false negative. Thus we investigate the structure of each component of category (iii) as follows: We label(a) each PI connected to all GTIs as likely true positive; (b) each GTI connected to all PIs as likely true positive; (c) each PI connected to some GTIs as likely false positive; and (d) each GTI connected to some PIs as likely false negative.

In order to assess the impact of every step of the pipeline leading to a set of predicted isoforms, we also built graphs for perfect read clusters (i.e. clustering the reads simulated from each isoform together) in three different ways. For the first graph, we generated a position-by-position consensus sequence for each perfect cluster by aligning its reads to their isoform-of-origin *V* = *S_T_* ∪*S*. Inaccuracies in this graph reflect only the sequencing error model. For the second graph, we generated a position-by-position consensus for each cluster but using the reads split-alignment to the genome rather than their isoform. This graph reflects the combined inaccuracies resulting from sequencing errors as well as split-alignment errors. Finally, for the third graph, we replace the clustering stage of Freddie with perfect clustering done manually. This graph reflects the error introduced by the segmentation stage in Freddie independently of its clustering stage. Fig. 6 plots various statistics on the structure of the resulting graphs for various threshold *t* values defining edges of the graphs.

**Fig. 6.**
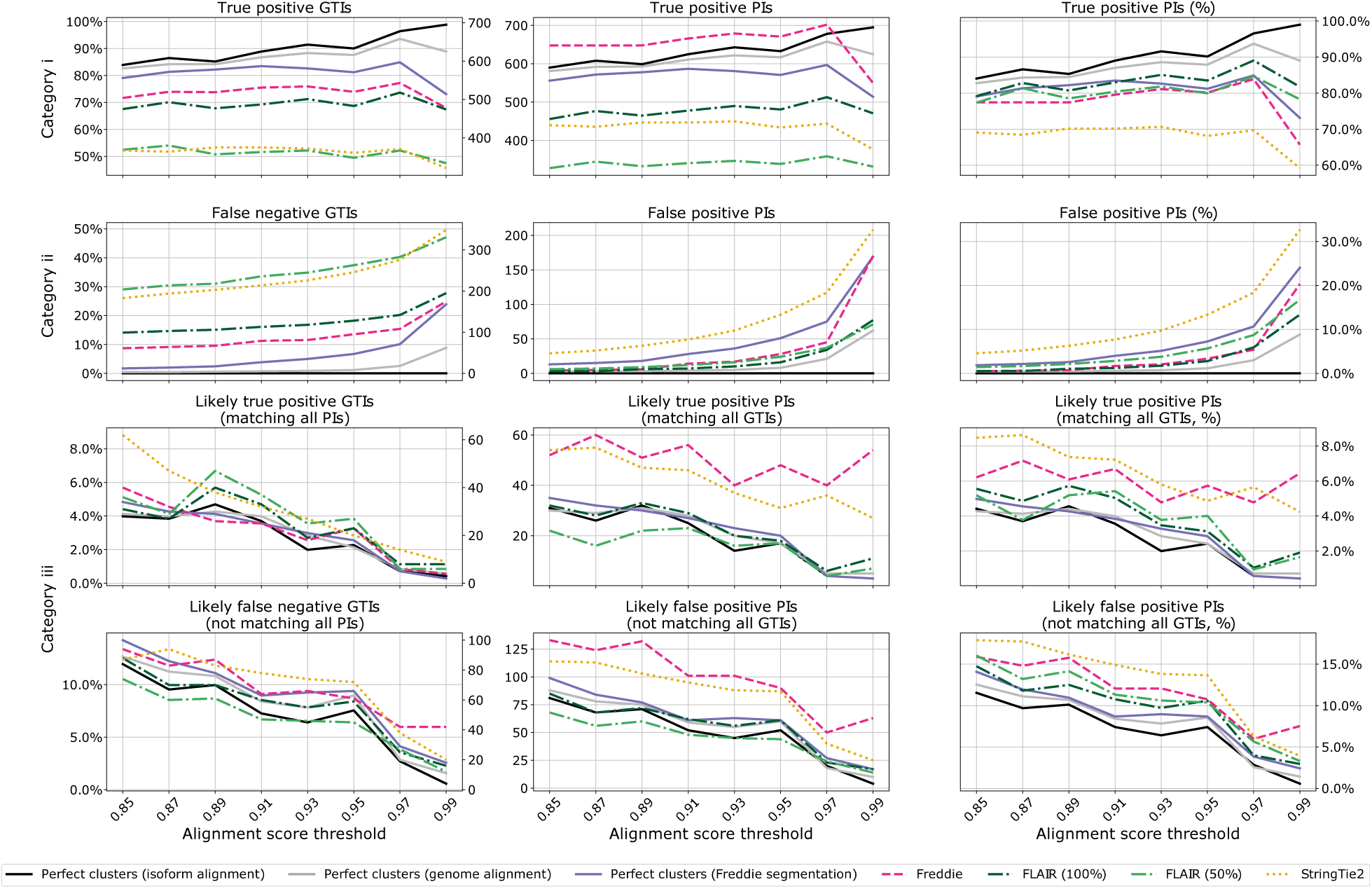
Graph-based accuracy results of the simulated dataset experiment. The first and second rows plot the count/percentages of isoforms belonging to mixed clique components (Category i) and non-mixed components (Category ii). The last two rows describe mixed ambiguous components (Category ii). The left column presents the absolute counts and percentages for GTIs in each type of components for all graphs. The middle and right columns present the absolute and percentage for PIs in each type of components for all graphs.

#### Simulated dataset results

While the assessment of the six graphs (three from the tool outputs and three from the perfect clustering), is fairly consistent across all threshold *t* values, we use *t* = 0.97 to quantify our findings since this value presents a clear inflection point in the plots. Freddie, FLAIR, and StringTie2 respectively predicted 837, 576, and 637 isoforms versus a total of 702 GTIs. The analysis of Fig. 6 shows that, overall, Freddie and FLAIR perform similarly, in terms of accuracy of the PIs compared to the GTIs; both tools outperform StringTie2 in terms of true positive and false negative, both in terms of absolute count and of proportion. We note that FLAIR’s results are obtained with the exact annotation for the GTIs in the dataset, while StringTie2 and Freddie are annotation-independent tools. If we run FLAIR with partial annotation dataset, we notice precipitous fall in true positive GTIs (see FLAIR %50 in Figure 6 and Figure S4)) The main difference between Freddie and FLAIR is that Freddie predicts significantly more isoforms; Figure 6 suggests that Freddie tends to separate reads from a given GTI into several clusters, which is not unexpected for a fully de-novo tool compared to an annotation-dependent tool such as FLAIR. However, in most cases, these clusters closely match the same set of GTIs, which is why we observe a higher raw true positive PI for Freddie even compared against the baseline graphs.

In terms of computational resources, StringTie2 had by far the smallest computational footprint, finishing in under a minute and with less than 26MB of RAM. Freddie and FLAIR had relatively more comparable RAM use at 1.00GB and 0.51GB, respectively. FLAIR outperformed Freddie in CPU time use, finishing in under two minutes compared to Freddie finishing in 20 minutes. Note that roughly three quarters of Freddie’s CPU time is spent in Gurobi ILP solver. We also include in the supplementary material (Table S1), the computational resources footprint for the tested tools on a larger dataset. The larger dataset is awhole genome simulated dataset generated using a procedure similar to the on described above, but without the restriction of using only chromosome 21.

#### Assessing the resolution of exon boundaries

The alignment-based similarity assess provides us with a good assessment of the structure of the predicted isoforms and their ground-truth counterparts. However, it is also important to understand the accuracy of the exact locations of exon boundaries generated by the different tools. Therefore, we extracted the set genomic locations of the exon boundaries of the ground truth isoforms and of the predicted isoforms of each tool. For each tool, We counted for each exon boundary locus the number of ground truth exon boundaries it has as a neighbor in a neighborhood of ±10bp. Figure 7 illustrates a histogram of these counts for the tested tools. As expected, we observe that FLAIR has the highest exon boundary resolution, with its histogram tightly concentrated at position zero. Freddie and StringTie2 have similar distributions with the absolute counts being higher for Freddie (Freddie predicts more isoforms). Generally speaking, all tools have histograms tightly concentrated around the zero position, indicated high base-level resolution across all tools.

**Fig. 7.**
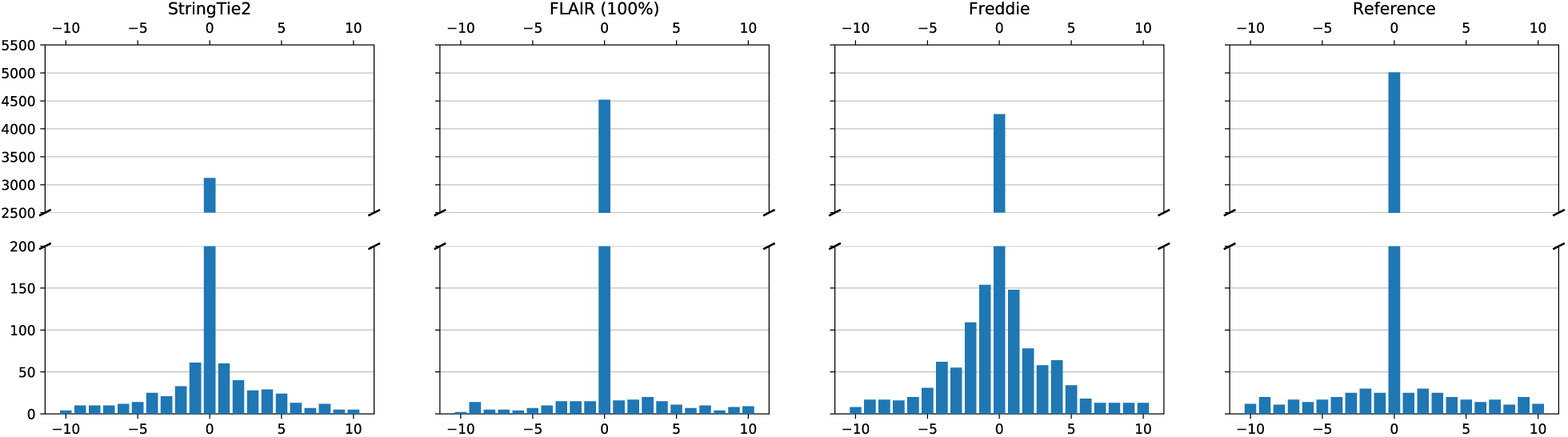
Histogram of the number of ground truth exon boundaries in the neighborhood of the predicted exon boundaries for different tools. As a baseline, we also show the histogram for the ground truth plotted against itself. For the baseline, bars not on the zero position are explained by the fact that some different annotation exons have very close starting/ending positions.

### 3.2 Real data experiment

#### Data generation

We generated a real mRNA transcriptomic dataset using Oxford Nanopore Technologies MinION PCR-free cDNA sequencing platform (chemistry kit SQK-DCS108). The mRNA material was extracted from a castration-resistant prostate cancer cell line called 22Rv1 [29]. This cell line is widely used as a first step model for studying prostate cancer. Over the past few years, a number of studies identified a number of alternatively spliced isoforms for the androgen receptor (AR) gene with potential links to the development of prostate cancer [10, 20]. Therefore, 22Rv1 and AR are promising subjects for discovering novel AS isoforms. In total, we generated 3.3 million long reads, with an average read length of 1587 nucleotides. In this experiment, we focused our analysis on the AR gene. Out of the 3.3 million reads, 839 mapped to the AR gene genomic region.

#### Detection of a potentially novel AR isoform

We ran Freddie on the AR reads we extracted from the real 22Rv1 LR dataset. We manually examined the predicted isoforms and compared them against the previously annotated AR isoforms by the ENSEMBL database and augmented it with extra information from [20]. We identified one potentially novel AR isoform with high read support. The novel isoform shares a similar structure to AR-204 isoform variant but with longer first and fourth exons. The novel isoform structure is illustrated in Figure 8.

**Fig. 8.**
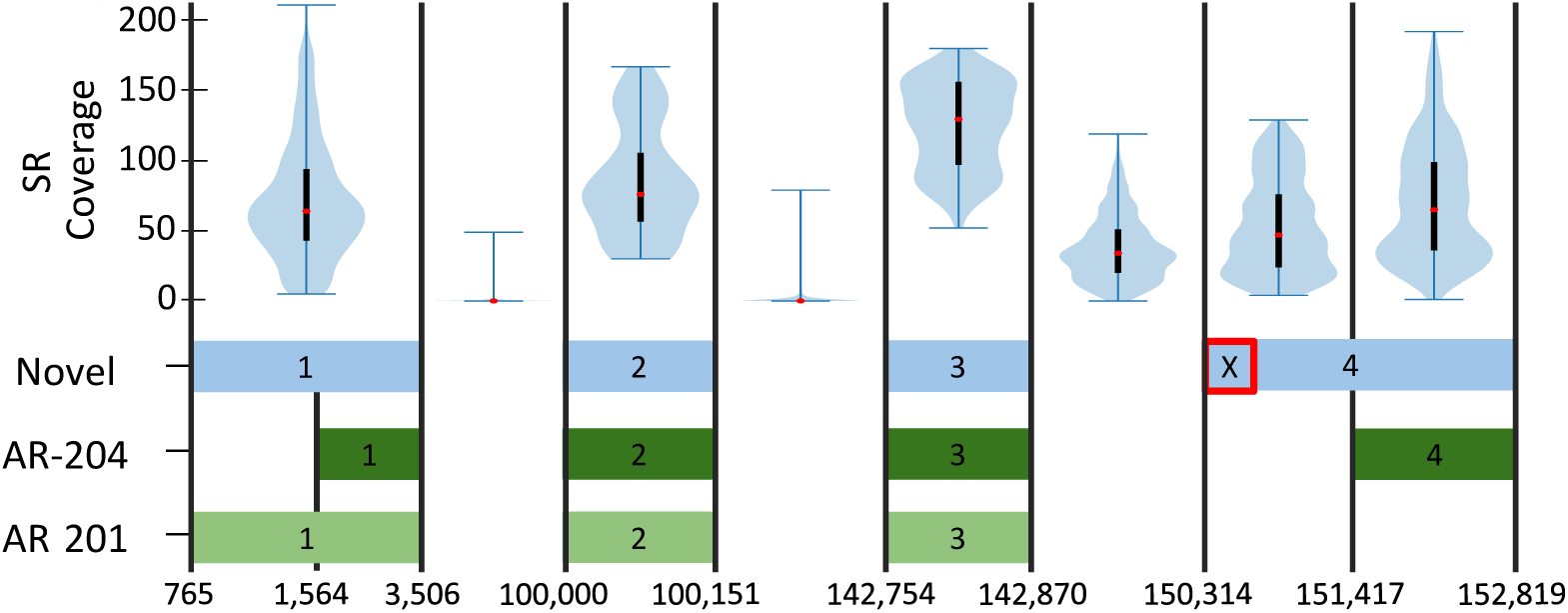
Potentially novel AR isoform detected by Freddie (Novel track). We show two annotated isoforms, AR-201 and AR-204, for AR as described in ENSEMBL database that have some shared exons with this novel isoform. We also annotate the location of “cryptic” exon #5 as described by Lu et al [20] by the X mark. The x-axis shows the genomic coordinated upstream of AR start location at chrX:67,543,256. The top track shows the short-read dataset coverage over the different exons and introns of AR using violin plots. The first and third quarterlies of the coverage distribution are indicated with the black boxes and its median is indicated with a red dot.

#### Orthogonal short-read validation

Since the new isoform we detect is not reported in ENSEMBL [3] or RefSeq [23] databases, we wanted to see if we can provide evidence for its structure using an orthogonal technology such as SR sequencing. As mentioned earlier, short reads are limited in their capacity to identify complete splicing structures of novel isoforms. However, in the case of a novel splice junction or an exon expansion into an intronic region, they can used to provide supporting evidence in a targeted manner.

Therefore, we downloaded a publicly accessible mRNA Illumina single-end short-read dataset for 22Rv1 (SRA accession number: SRR8446406) [27]. Since the novel detected isoform includes a genomic segment that is specific to it and not covered by the exons of any of the other annotated isoforms, we wanted to see if mapping the SRs to the reference genome results in coverage of this specific genomic segment which is intronic for all annotated isoforms. We used BWA-MEM [14] to map the SR read dataset to the human reference genome. The region of the partially retained intron 3 has coverage median of 47x, significantly higher than intron 1 and 2 of AR. The non-retained part of intron 3 has coverage of 34x potentially as a result of cryptic exons belonging to this region [20]. Figure 8 provides a visualization of the genomic alignment coverage of the SRs on this specific genomic segment.

## 4 Discussion and conclusion

Freddie is a novel alternative splicing isoform detection tool that does not rely on isoform annotation databases. Freddie is designed to address the characteristic sequencing errors of LRs, especially in the absence of isoform annotations. Using simulated data, we demonstrate that Freddie achieves better recall than existing detection tools even when they are supplied with isoform annotations. Freddie also has a similar false positive rate to the annotation-dependent tool FLAIR. On real dataset, we used Freddie to detect a potentially novel androgen receptor isoform and used an orthogonal short-read dataset to validate it. While this is promising evidence, we believe that further validation of this isoform is warranted.

As future directions, we plan to further analyze the computational complexity of the MErCI problem used in Freddie. We hope to identify a tight bound on the complexity of solving MErCI and shed light on possible ways to speed it up. We will also aim to create variants of MErCI to address the problem of detection of other transcriptomic targets, besides AS isoforms, such as circular RNA and gene fusions.

## Acknowledgement

We would like to thank Dr. Zoubeidi and her team at the Vancouver Prostate Centre for providing us with 22Rv1 cell culture to perform MinION Sequencing.

