## Supplementary material for "Freddie: Annotation-independent Detection and Discovery of Transcriptomic Alternative Splicing Isoforms"

### Freddie (Supplementary Material)

Orabi et al.

January 20, 2021

Table 1: Computational resources use on whole transcriptome sequencing. CPU (user) time, real (clock) time, and memory use of FLAIR and the different stages of Freddie. Note that we do not report the results for StringTie2 because it crashes when run on the whole genome sample (segfault).

| Tool/stage | Thread count | CPU time (min) | Real time (min) | Max memory use (GB) |
|---|---|---|---|---|
| FLAIR | 32 | 350.43 | 21.42 | 11.64 |
| Freddie split | 1 | 79.18 | 82.75 | 3.31 |
| Freddie segment | 32 | 567.69 | 47.66 | 3.15 |
| Freddie cluster | 32 | 2299.17 | 106.97 | 9.80 |

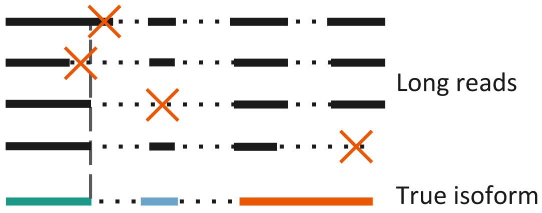


Figure S1: Long-read sequencing has a high error rate reaching up to 15%. Insertion and deletion (indel) of nucleotides dominate this error rate. These indel errors results in 1) a reduction in the resolution of exon boundaries and 2) missing small exons. Additionally, 3) full-length LR sequencing is not always successful, resulting in missing exons at the tail of the sequenced isoform. The figure illustrate these three hazards associated with LR sequencing of isoforms.

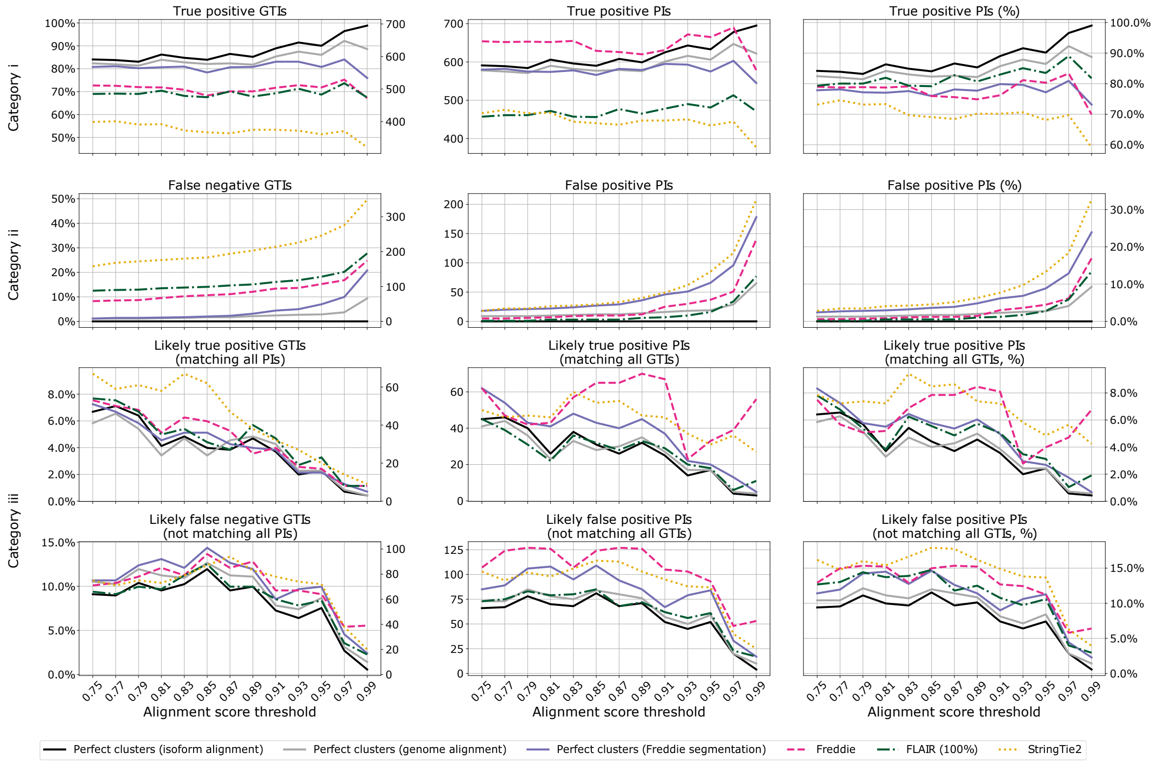


Figure S2: Graph-based analysis with Minimap2 mapper used for all tools.

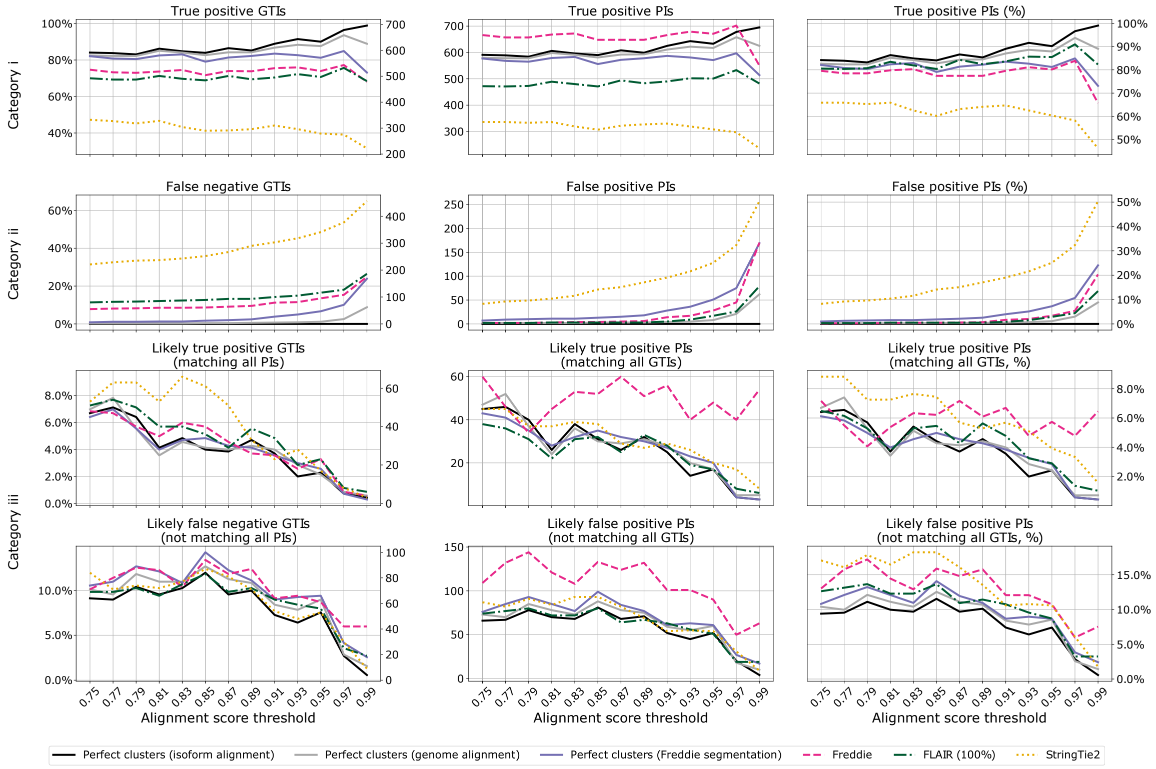


Figure S3: Graph-based analysis with deSALT mapper used for all tools.

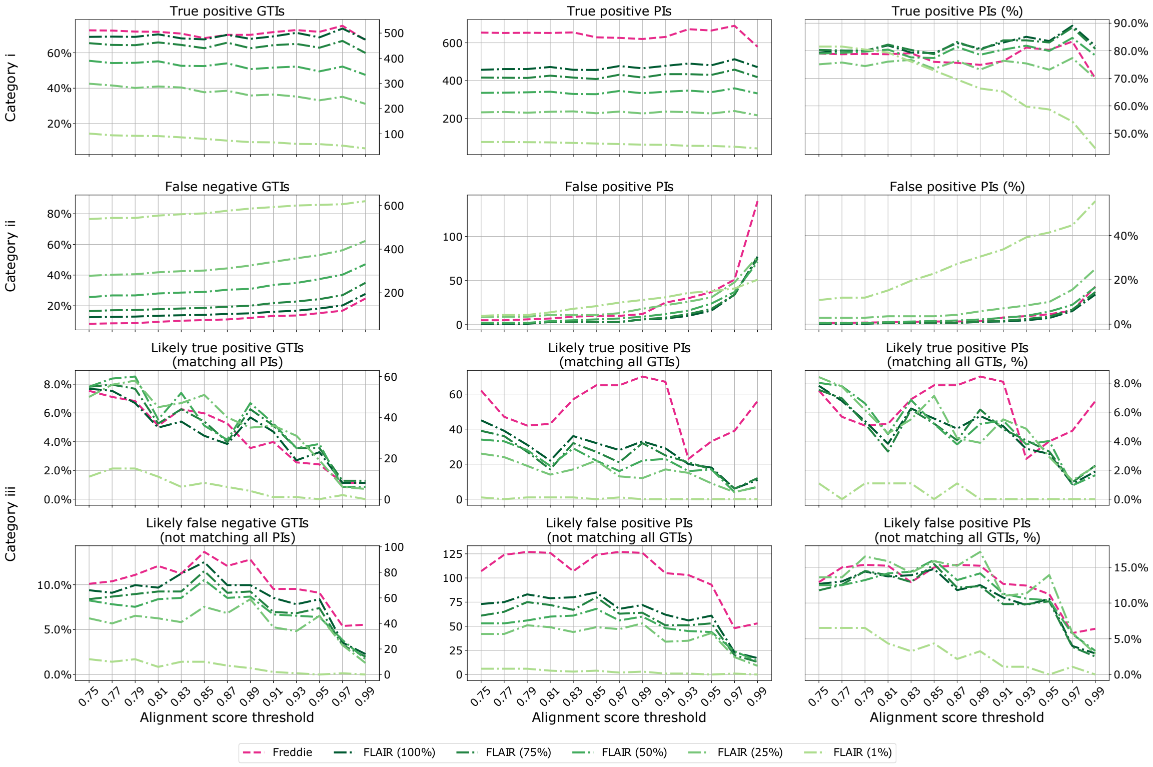


Figure S4: Graph-based analysis of Freddie vs FLAIR with Minimap2 mapper when FLAIR is given varying sample sizes of chromosome 21 annotations.
